## Supplementary Material for "Structure-guided glyco-engineering of ACE2 for improved potency as soluble SARS-CoV-2 decoy receptor"

†equal contributions.

*Correspondence to:

**Table S1.** Glycosylation sites and N-glycan structures used in the models of SARS-CoV-2 Spike and human ACE2 are indicated. Blue squares indicate N-acetylglucosamine, green circles mannose, yellow circles galactose, purple squares sialic acid and green triangles fucose residues. Anomericity and positions of glycosidic linkages are indicated where necessary.

| Protein | Glycosylation site | Glycan structure |
| --- | --- | --- |
| SARS-CoV-2 Spike | N61 | 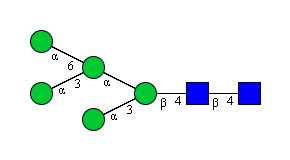 |
|  | N74  N122  N149  N165 | 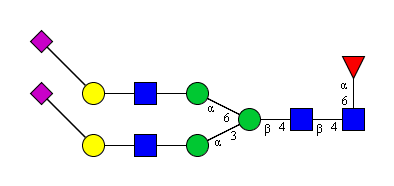 |
|  | N234 | 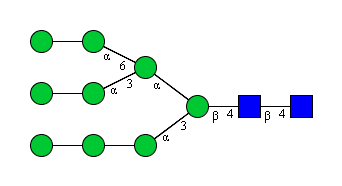 |
|  | N282  N331  N343 | 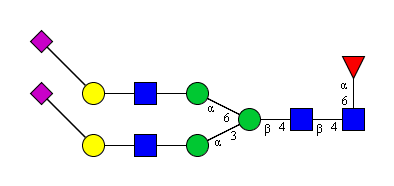 |
|  | N603 | 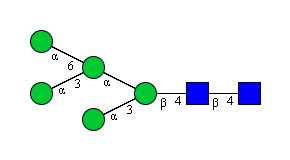 |
|  | N616  N657 | 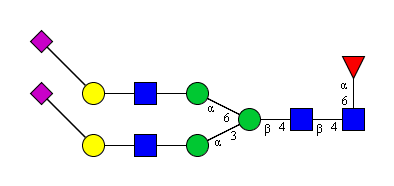 |
|  | N709  N717  N801 | 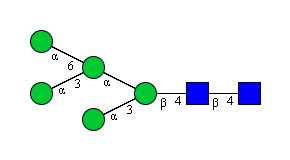 |
|  | N1098  N1134 | 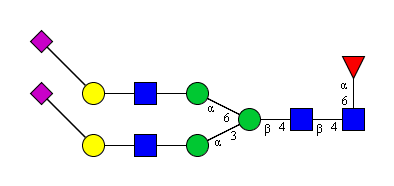 |
| human ACE2 | N53  N90  N103  N322  N432  N546  N690 | 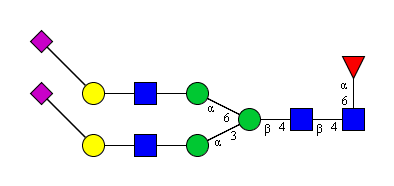 |

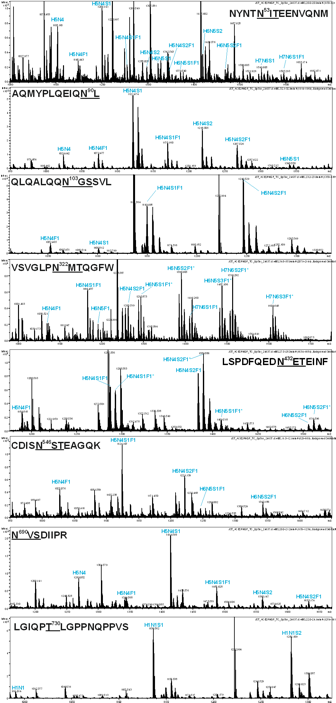

**Figure S1. Site-specific glycosylation profiles of rshACE2.** Prior to analysis by LC-ESI-MS, reduced and S-alkylated rshACE2 was digested in-solution with chymotrypsin and trypsin. Glycan compositions are denoted as the sum of hexoses (H), N-acetyl-hexosamines (N), N-acetyl-neuraminic acids (i.e. sialic acids; S) and fucoses (F). Molecular ion-species detected as ammonium adducts (i.e. +17 amu) are indicated by asterisks.

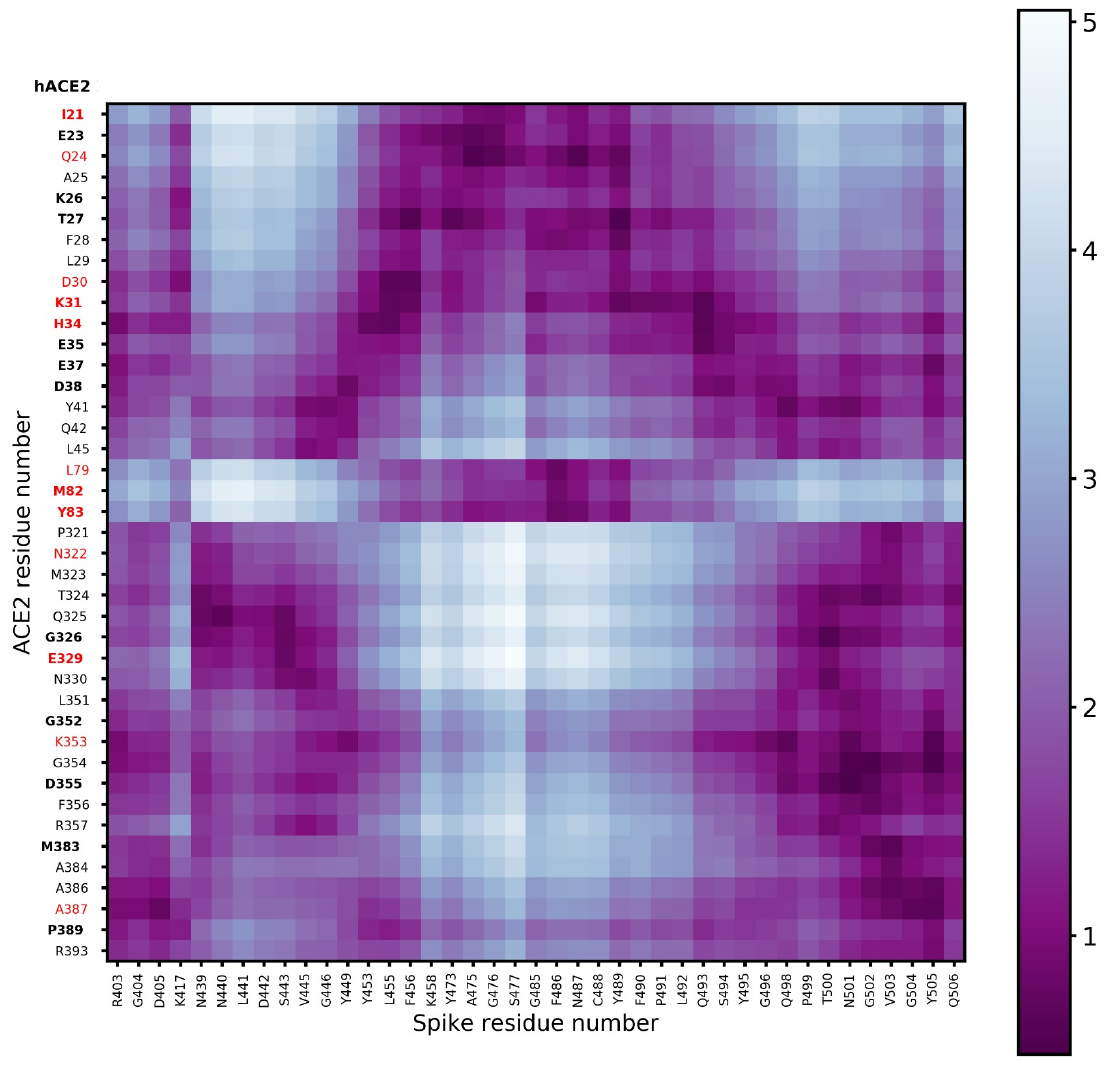

**Figure S2.** **Distance heatmap of human ACE2 (hACE2) residues in contact with Spike RBD**. Black squares correspond to an average distance of 0 nm and white squares to an average distance of 6 nm. Residues which differ in mouse ACE2 are highlighted in red, residues known to be genetically polymorphic^[1]^ are shown in bold.

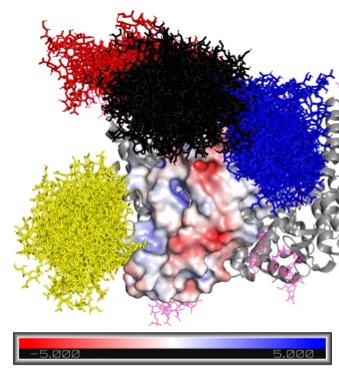

**Figure S3.** **Electrostatic potential of the binding interface region on ACE2.** Electrostatic potential surface of human ACE2. Red corresponds to a negative charge and blue to a positive charge. A bundle of glycan conformations is shown in sticks for the glycans at N53 (blue), N90 (yellow), N322 (black) and N546 (red).

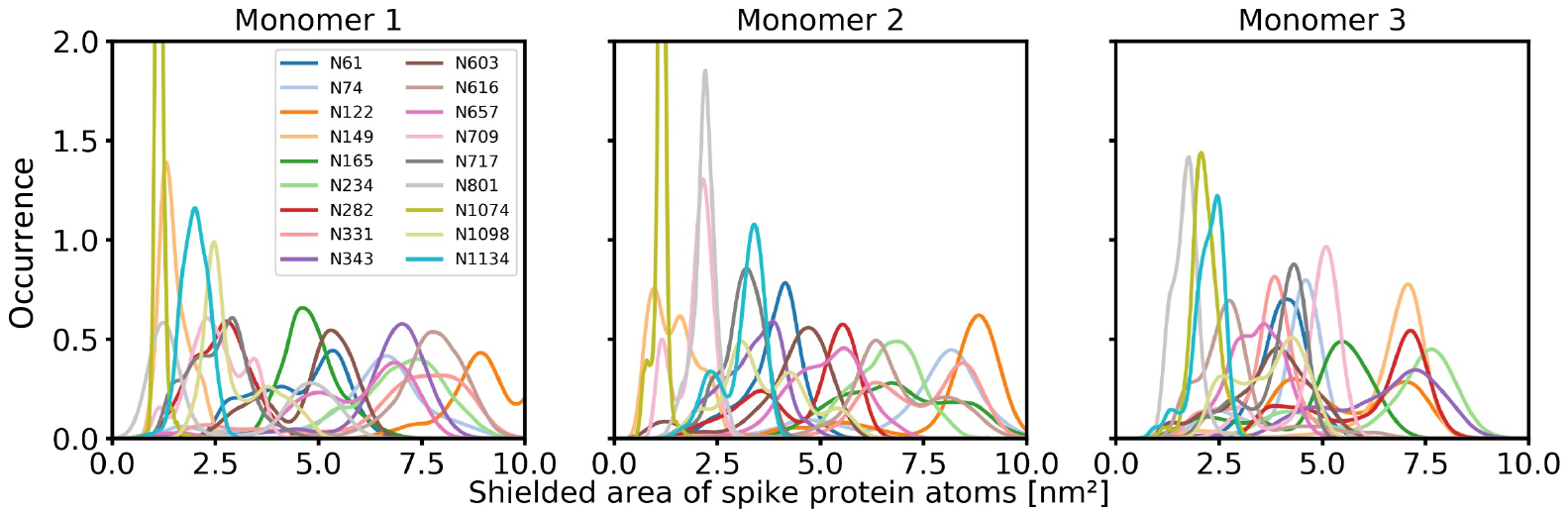

**Figure S4.** **Normalized distribution of the area of Spike atoms shielded by each of its N-glycans.** The distributions were obtained from the molecular dynamics simulations for the individual monomers of Spike. For every Spike glycosite, the area of Spike protein atoms that is shielded by the glycan during the simulation is indicated. For instance, the glycan at N331 on monomer 1 shields between 7.5 and 10.0 nm^2^ of the Spike surface (see also Fig. 1c of the main manuscript).

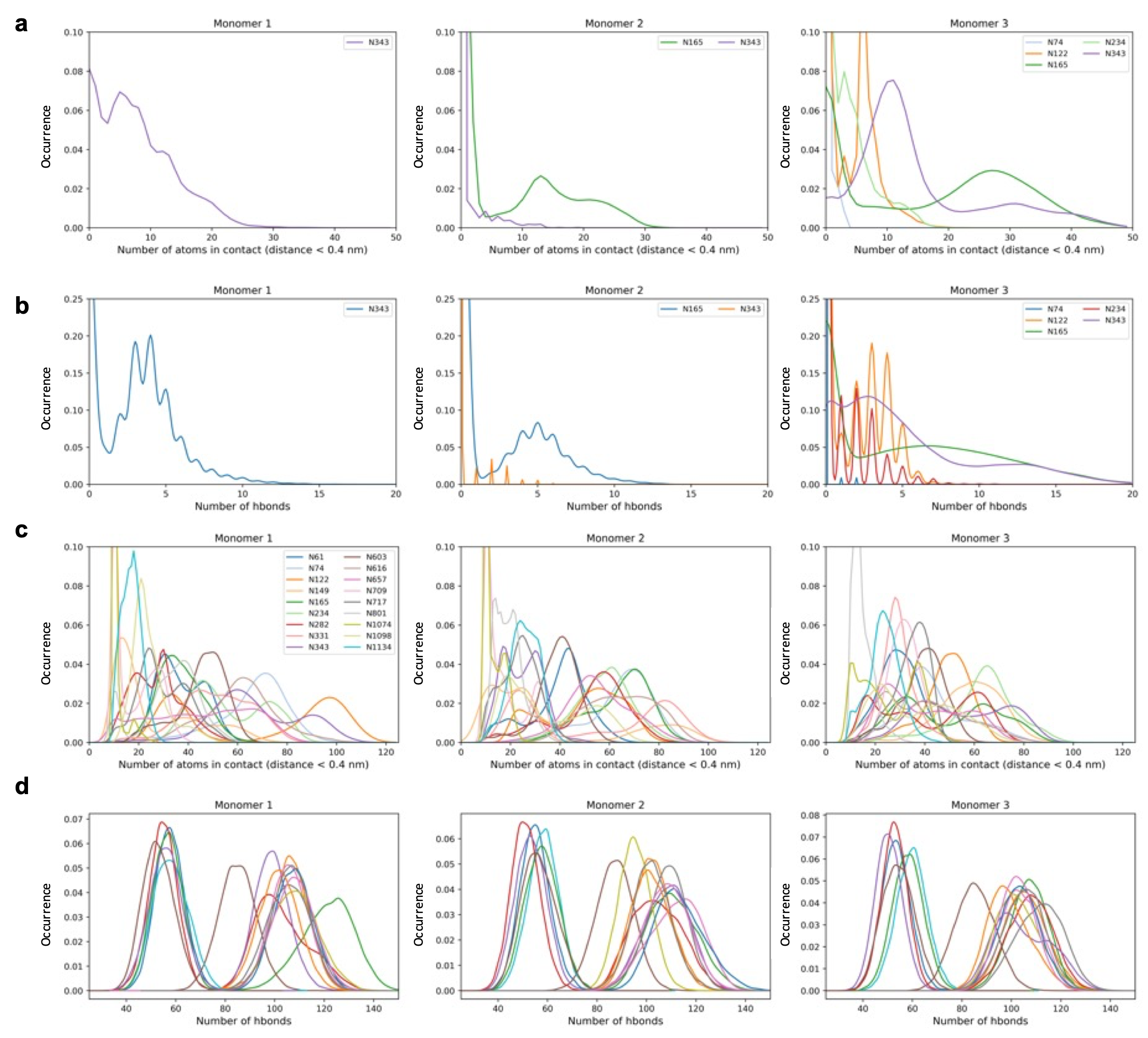

**Figure S5. Interactions of Spike N-glycans with ACE2 and with Spike. (a)** Normalized distribution of the number of atoms of glycans of the indicated Spike monomers that are in contact with human ACE2. **(b)** Normalized distribution of the number of hydrogen bonds formed between Spike glycans and human ACE2, shown for 2 Spike monomers. Glycans of the second monomer did not form H-bonds with human ACE2. **(c)** Normalized distribution of the number of atoms of Spike glycans that are in contact with Spike protein atoms, shown for all 3 Spike monomers. **(d)** Normalized distribution of the number of hydrogen bonds formed between Spike glycans and Spike for all 3 Spike monomers. Monomer 3 is the interacting monomer containing the RBD. The number of interacting atoms and the number of hydrogen bonds observed during the MD simulations are represented as normalized distribution for the individual glycosites on Spike on the different monomers.

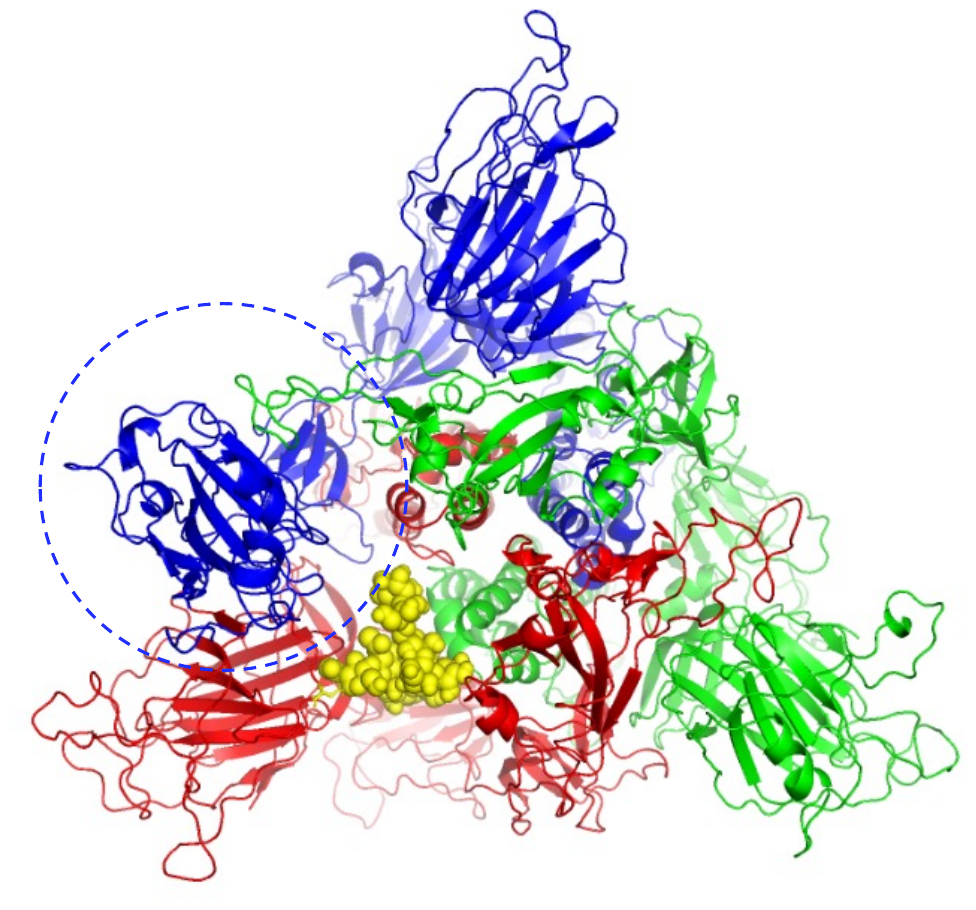

**Figure S6.** **3D** **positional modelling of glycan N234 in the Spike trimer.** The glycan at N234 (yellow spheres) of monomer 2 (red) is partially inserting into the core of the Spike trimer. The vacant space in the core is created by the receptor-binding domain of monomer 3 (blue) residing in the up conformation (in the dashed circle). Monomer 1 is shown in green.

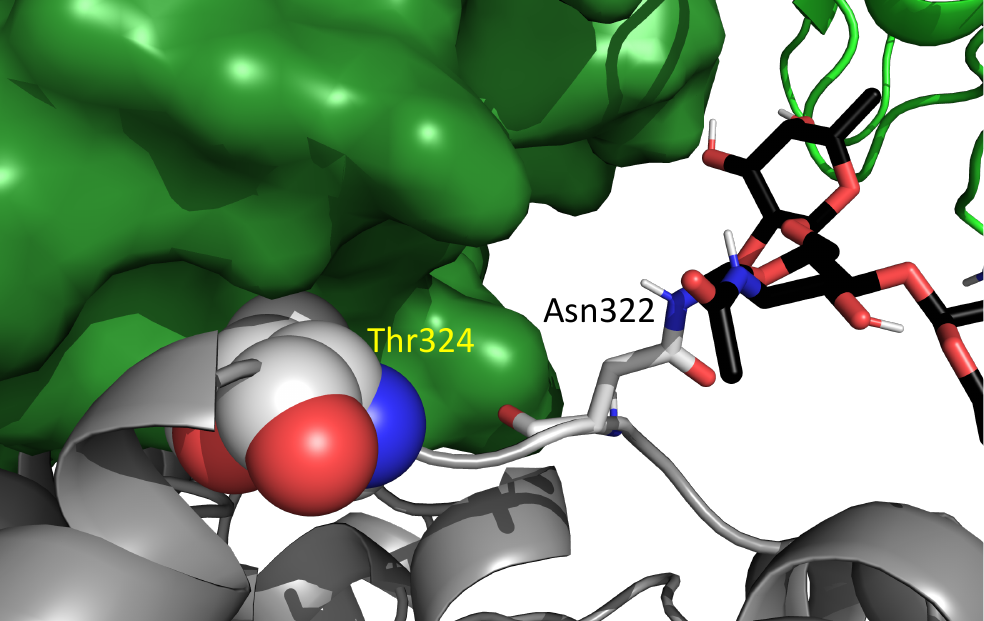

**Figure S7. Residue T324 of ACE2 at the interface to Spike.** Residue T324 (in spheres) of ACE2 (grey) is directly interacting with the RBD of Spike (dark green surface). Ablation of the glycan (black sticks) at N322 (grey sticks) was therefore achieved through the mutation N322Q.

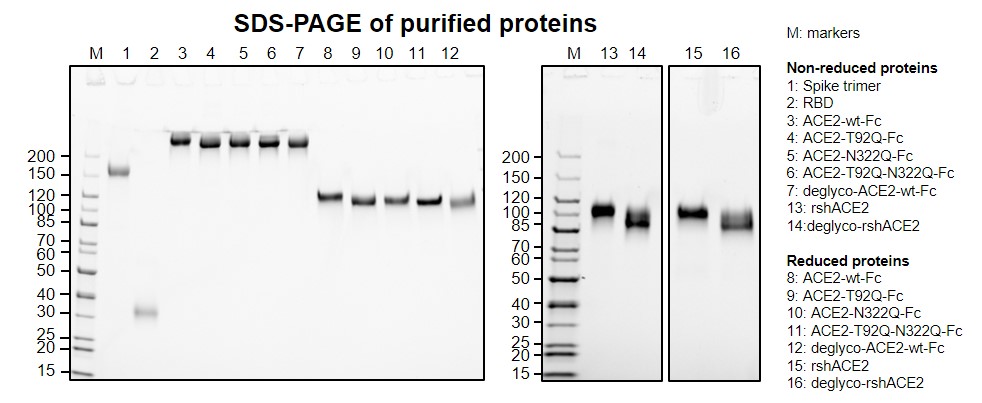

**Figure S8. Analysis of ACE2 and Spike variants by SDS-PAGE.** Analysis of 1 µg of each purified protein by SDS-PAGE. M, molecular mass markers.

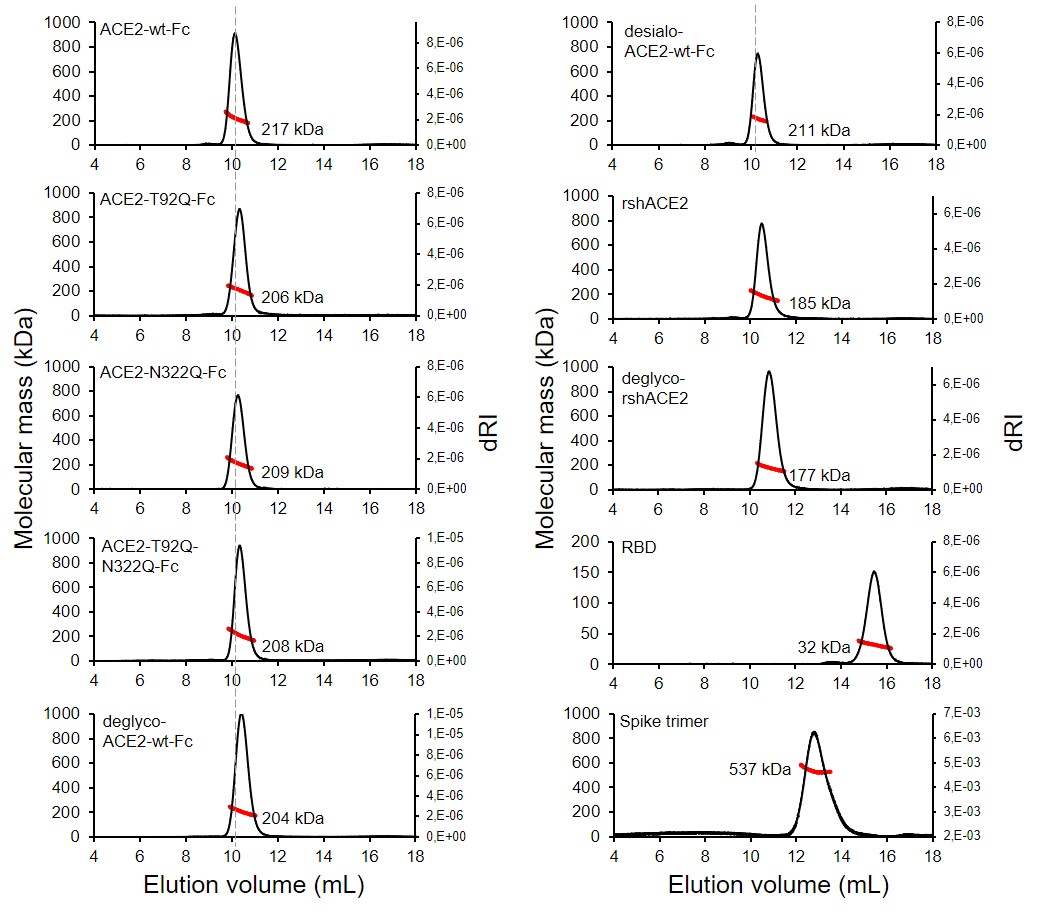

**Figure S9. Analysis of ACE2 and Spike variants by SEC-MALS.** A total of 50 µg of Spike was loaded onto a Superose 6 Increase 10/300 GL column at a flow rate of 0.25 mL min^-1^. All other proteins were analyzed by injection of a total of 25 µg of the respective protein onto a Superdex 200 10/300 GL column at a flow rate of 0.75 mL min^-1^. The elution time of ACE2-wt-Fc is marked with a dashed line. Molecular masses (red traces) were determined by MALS. dRI, change in refractive index.

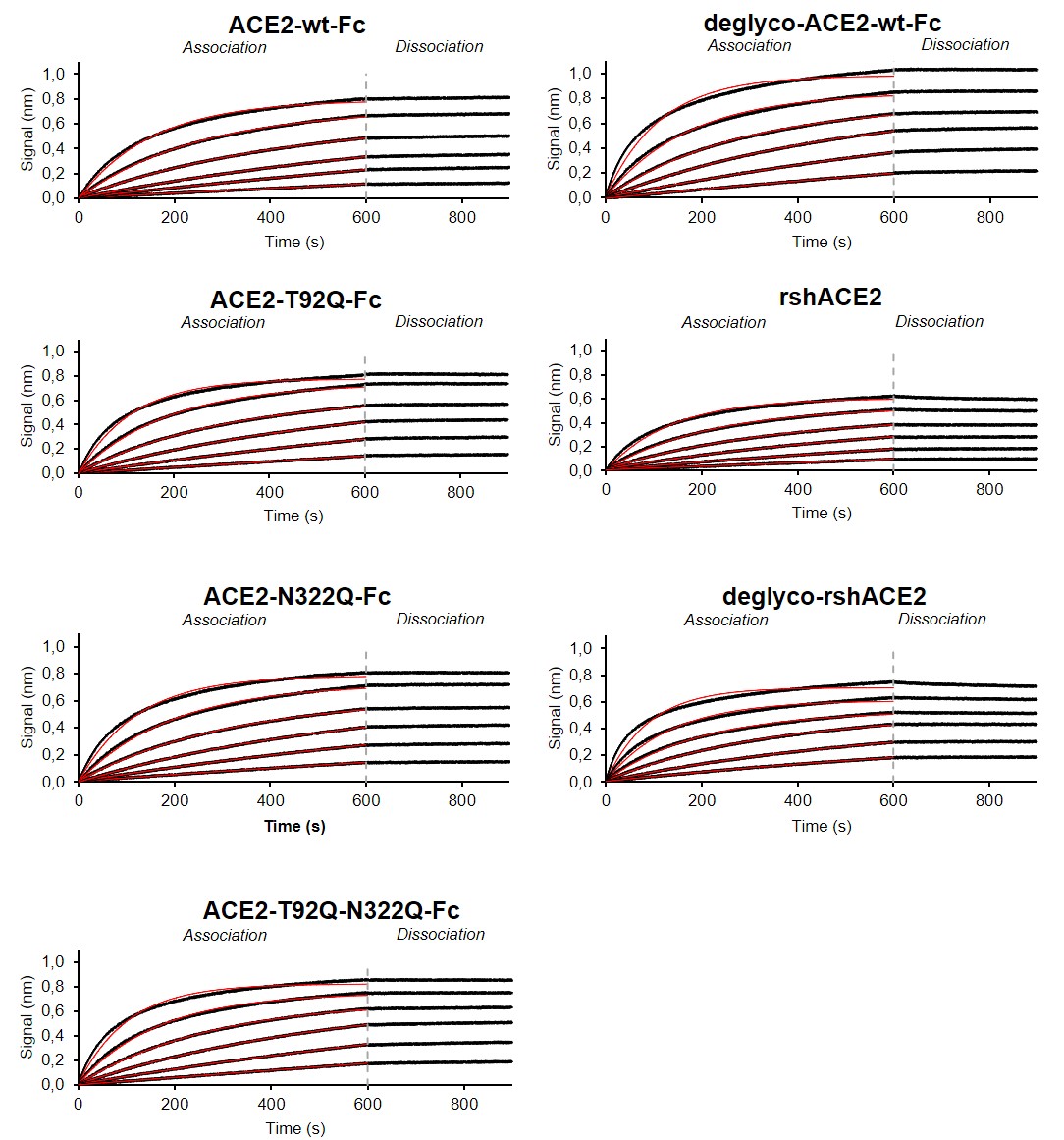

**Figure S10. Binding of Spike to human ACE2 variants as determined by biolayer interferometry.** Binding analysis was performed by dipping biosensors loaded with different ACE2 variants into 2-fold serial dilutions of purified Spike (1.6-50 nM). Black lines represent the response curves of the association and dissociation phases. The data were fitted to a 1:1 binding model (red lines). For each ACE2 variant one representative experiment out of three is shown.

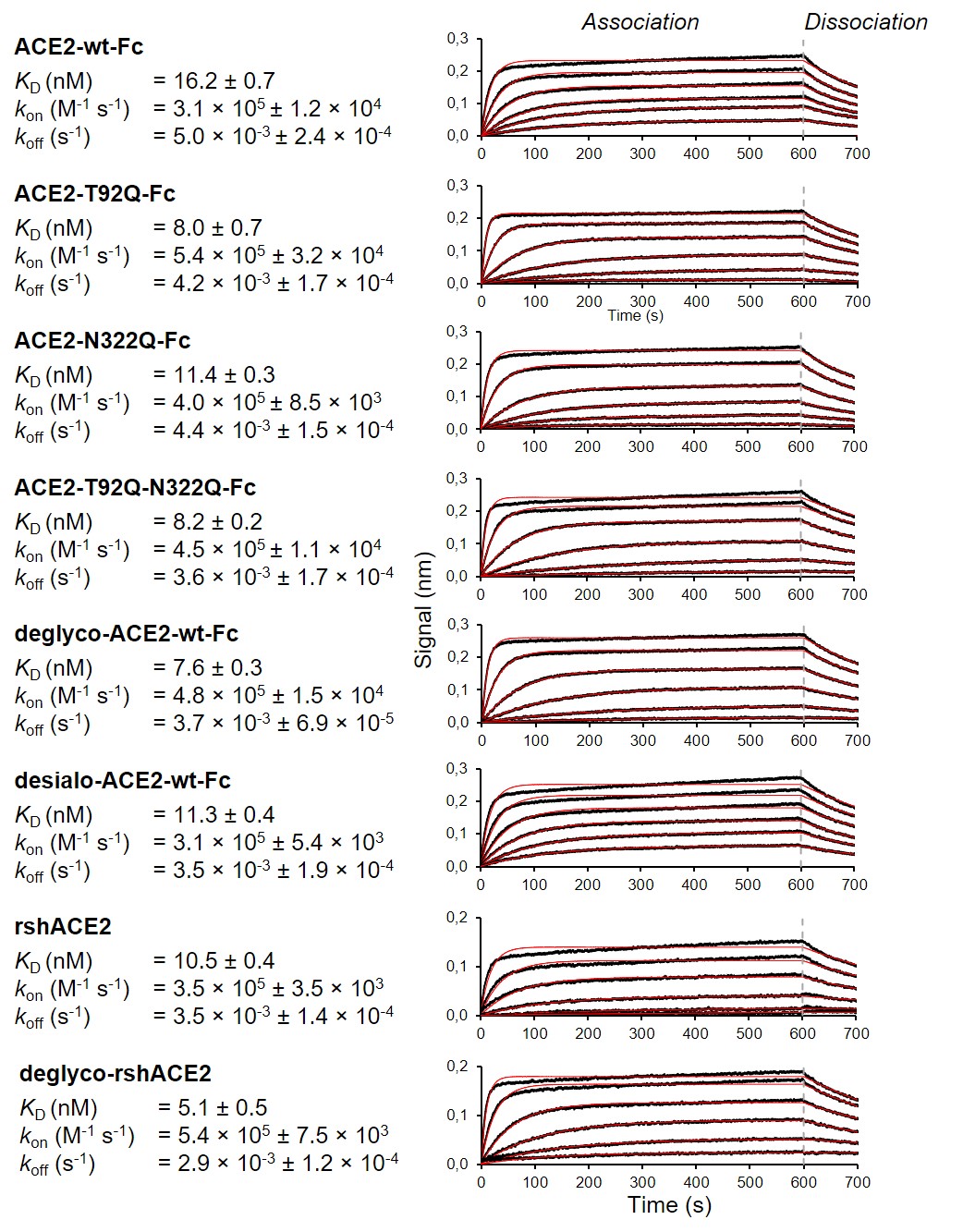

**Figure S11. Binding kinetics of RBD to human ACE2 variants as determined by biolayer interferometry.** Black lines represent the response curves of the association and dissociation phases. The data were fitted to a 1:1 binding model (red lines). *K*_D_, *k*_on_ and *k*_off_ values represent the mean ± SEM of 3 independent experiments. For each ACE2 variant one representative experiment is shown.

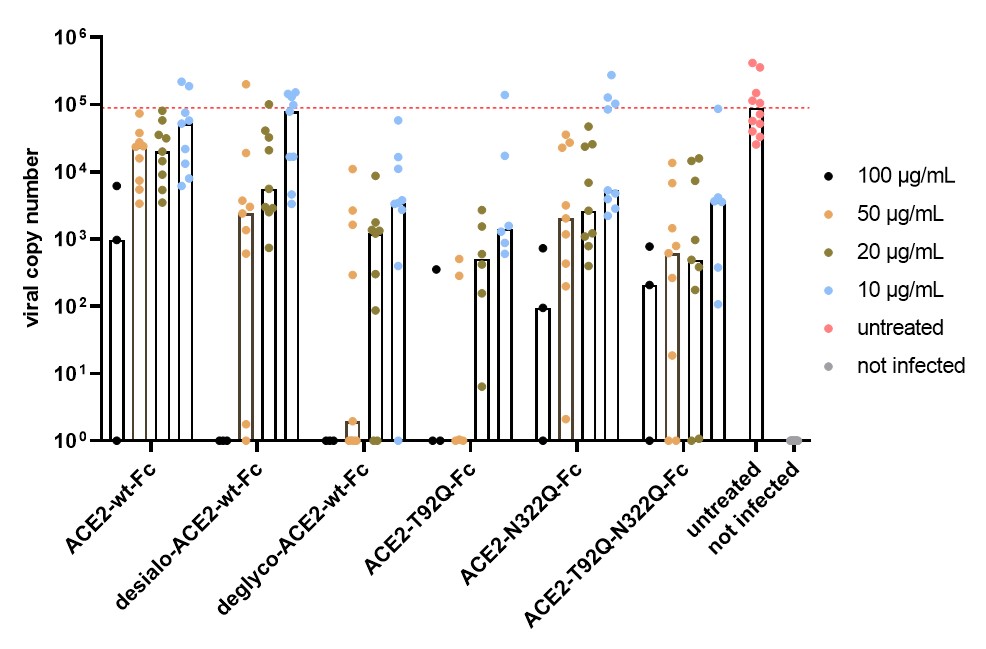

**Figure S12. Critical role of ACE2 glycosylation for SARS-CoV-2 infectivity.** Inhibition of SARS-CoV-2 infection of Vero E6 cells (MOI: 0.002) using wild-type ACE2-Fc and the indicated glyco-engineered ACE2-Fc variants at final concentrations of 10-100 µg/mL. The viral RNA content of the culture supernatants was quantified by RT-qPCR. Data are derived from 1-4 experiments performed in triplicates. Deglyco-ACE2-wt-Fc, enzymatically deglycosylated wild-type ACE2-Fc; desialo-ACE2-wt-Fc, enzymatically desialylated wild-type ACE2-Fc.

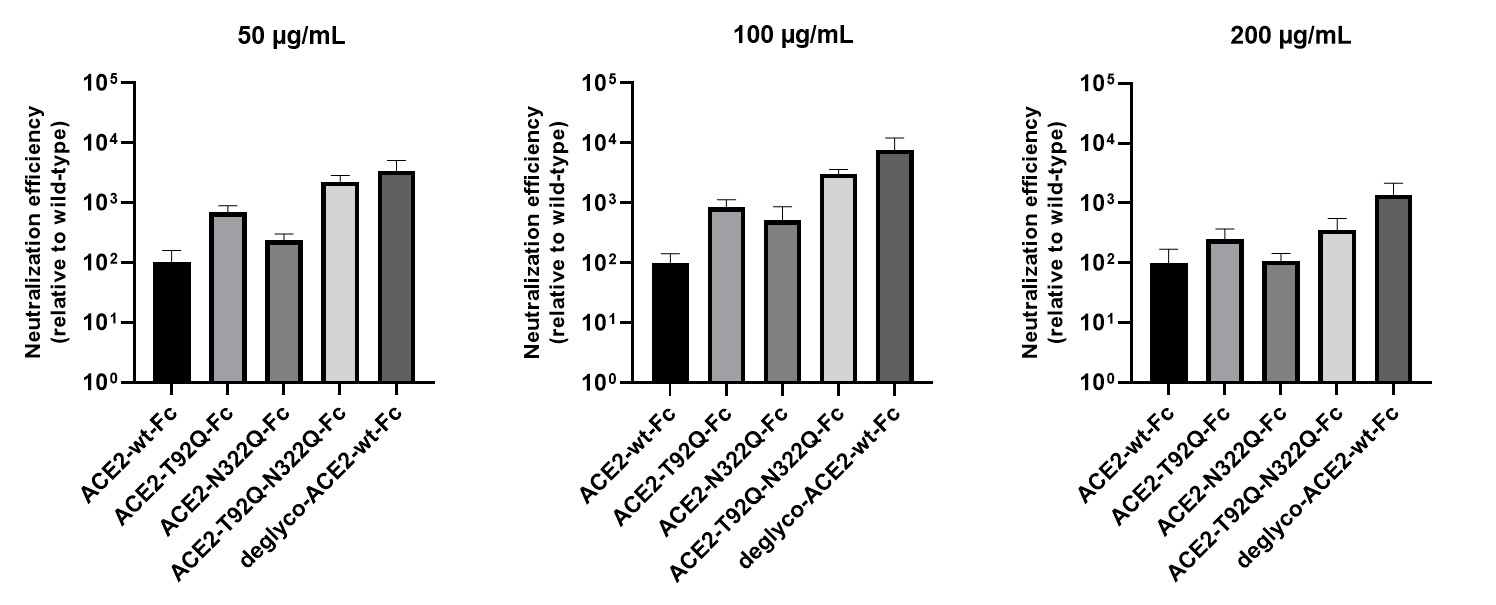

**Figure S13. Critical role of ACE2 glycosylation for SARS-CoV-2 infectivity.** Inhibition of SARS-CoV-2 infection of Vero E6 cells (MOI: 20) using wild-type ACE2-Fc and the indicated glyco-engineered ACE2-Fc variants at final concentrations of 50-200 µg/mL. The viral RNA content of the infected cells was quantified by RT-qPCR and expressed as neutralization efficiency relative to ACE2-wt-Fc (set to 100%). Data are presented as mean ± SD of triplicates. Deglyco-ACE2-wt-Fc, enzymatically deglycosylated wild-type ACE2-Fc.

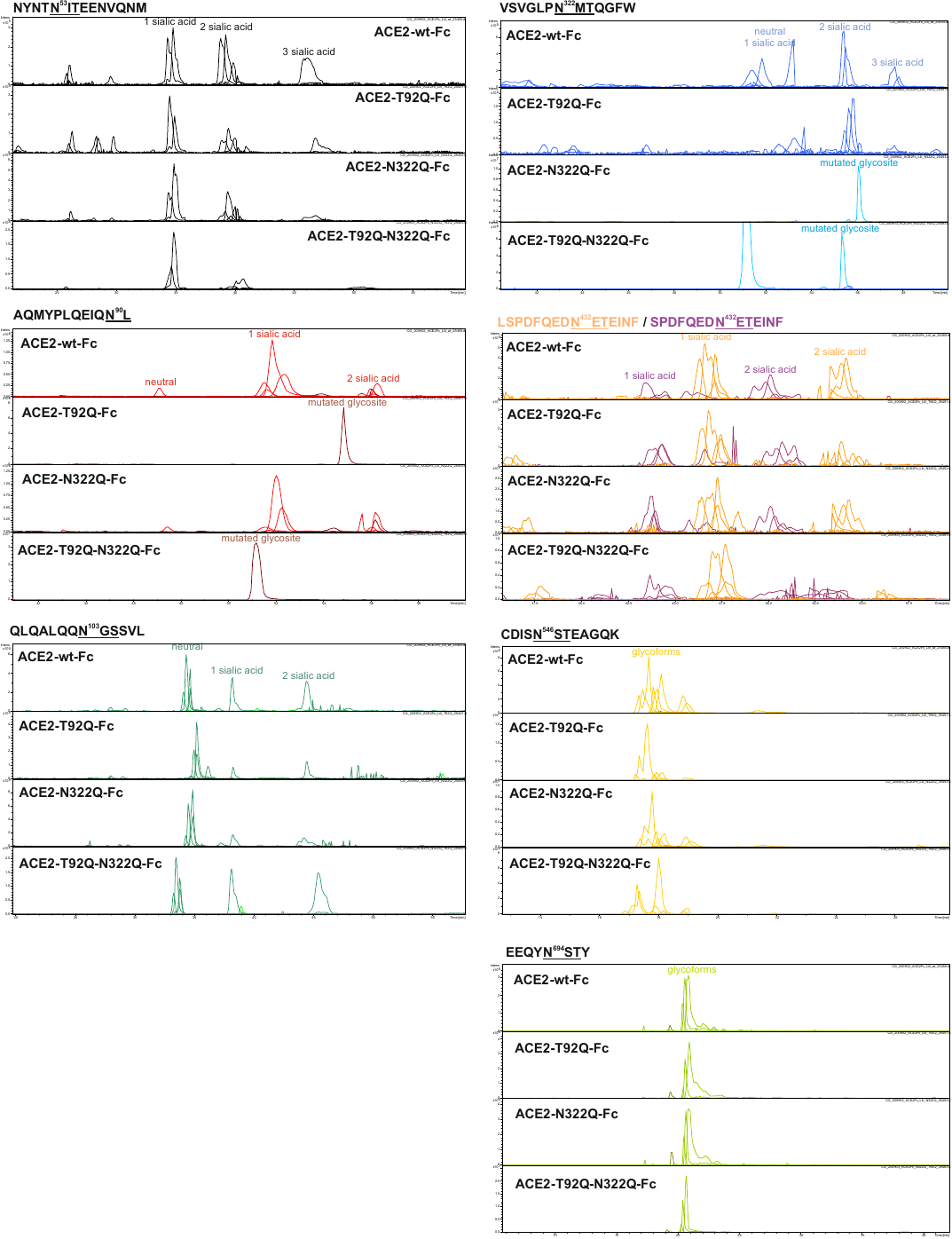
**Figure S14. Site-specific ablation of ACE2 N-glycans by mutagenesis.** Prior to analysis by LC-ESI-MS/MS, the indicated reduced and S-alkylated ACE2-Fc variants were digested in-solution with chymotrypsin and trypsin. Comparative analysis of extracted ion chromatograms of glycopeptides confirmed the absence of glycans attached to N90 and/or N322 in ACE2-T92Q-Fc, ACE2-N322Q-Fc and ACE2-T92Q-N322Q-Fc. The peptide sequences surrounding the different glycosites are shown.

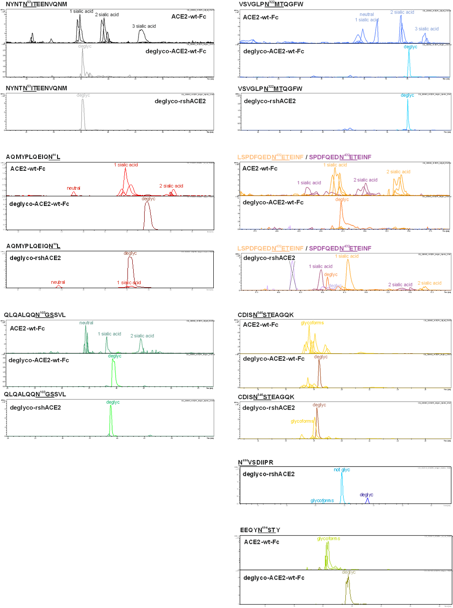
**Figure S15. Deglycosylation of ACE2-wt-Fc and rshACE2 by PNGase F.** Prior to analysis by LC-ESI-MS/MS, native and deglycosylated (deglyco) ACE2-wt-Fc and rshACE2 were reduced, alkylated and then digested in-solution with chymotrypsin and trypsin. Comparative analysis of extracted ion chromatograms of glycopeptides confirmed the absence of N-glycans attached to N53, N90, N103 and N322 of deglyco-ACE2-wt-Fc and deglyco-rshACE2. Peptide sequences surrounding the different glycosites are shown.

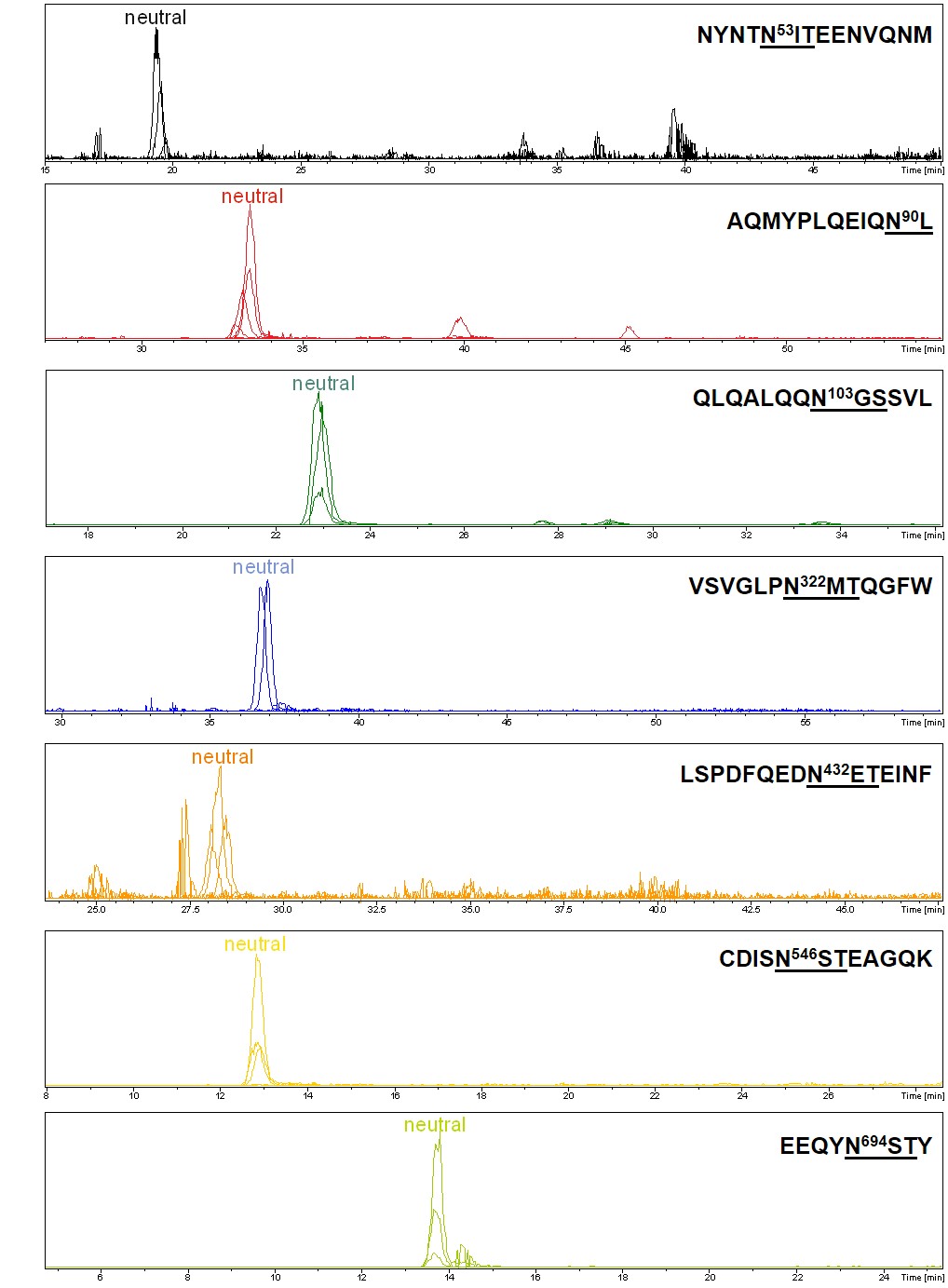

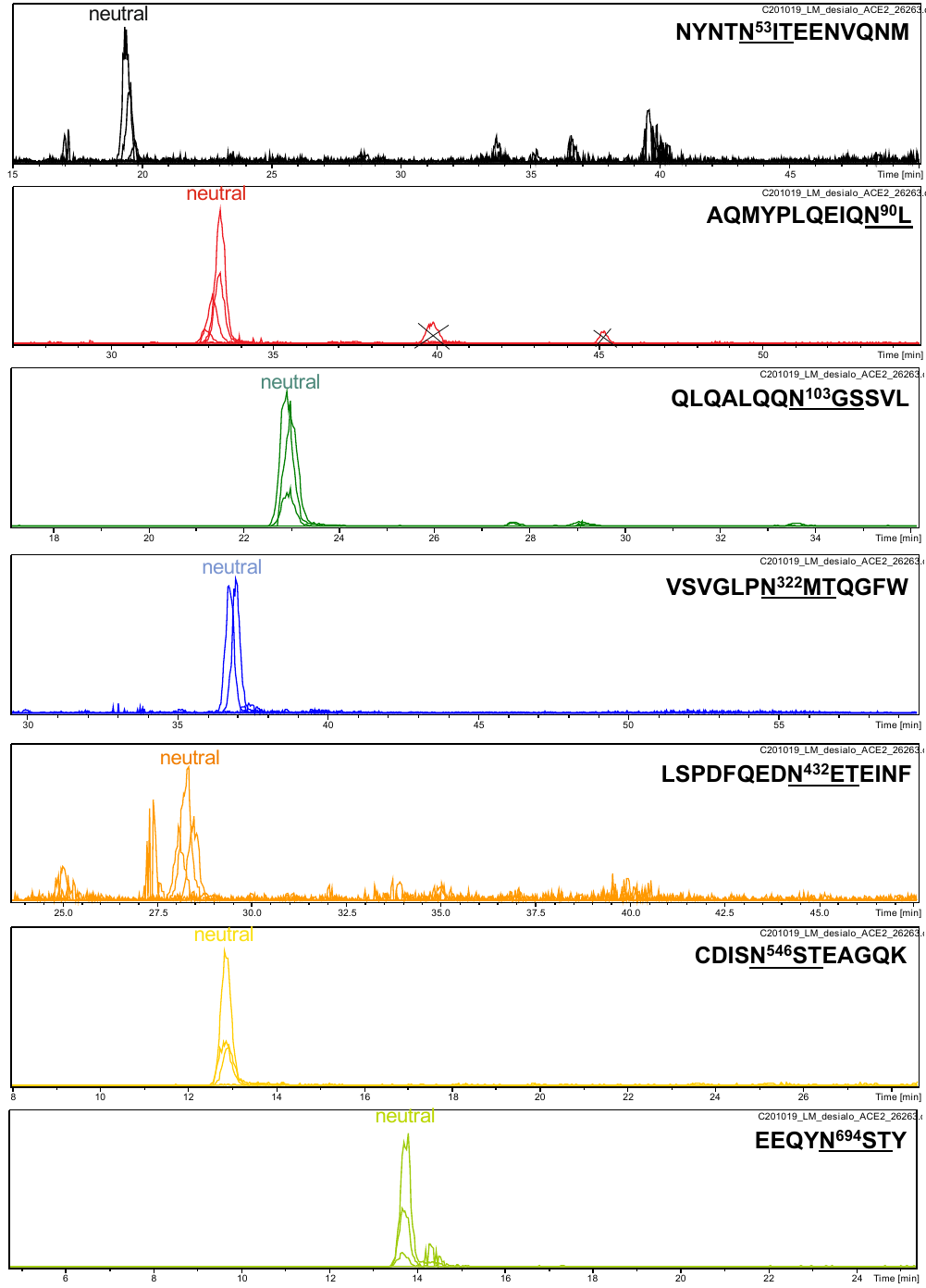
**Figure S16. Analysis of enzymatically desialylated ACE2-wt-Fc.** The absence of sialic acids attached to N53, N90, N103, N322, N432 and N546 of desialo-ACE2-wt-Fc was confirmed by ESI-LC-MS/MS. Peptide sequences surrounding the different glycosites are shown.

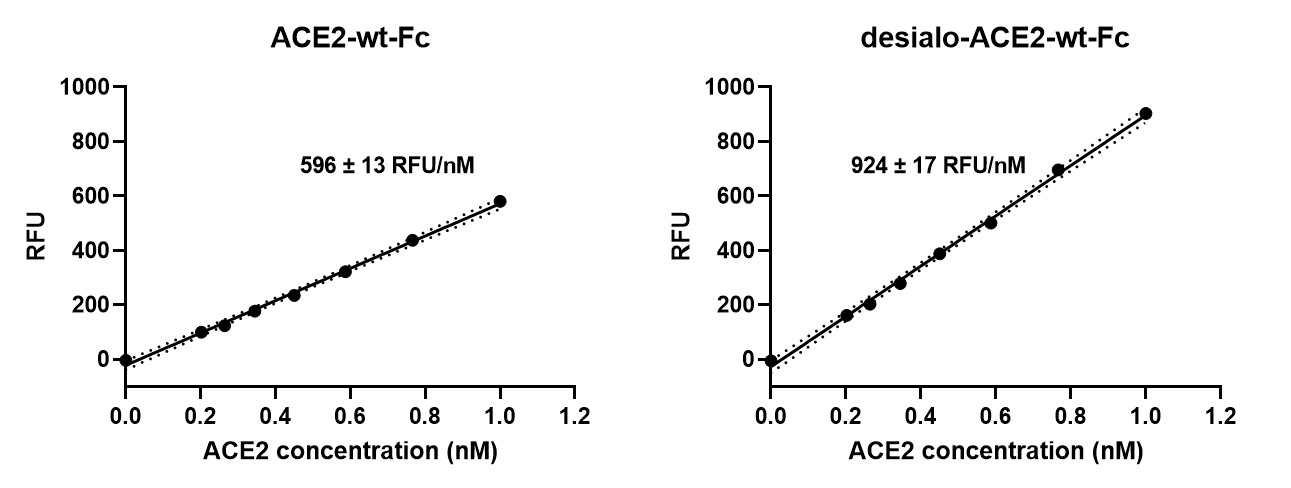

**Figure S17. Enzymatic activity of desialo-ACE2-wt-Fc.** Hydrolysis of 100 µM 7-methoxycoumarin-4-yl-acetyl-Ala-Pro-Lys-2,4-dinitrophenyl was continuously monitored by spectrofluorimetry. Hydrolytic activity is plotted as relative fluorescence units (RFU) over ACE2 concentration (in nM). All assays were performed in triplicates. One representative experiment out of three is shown.

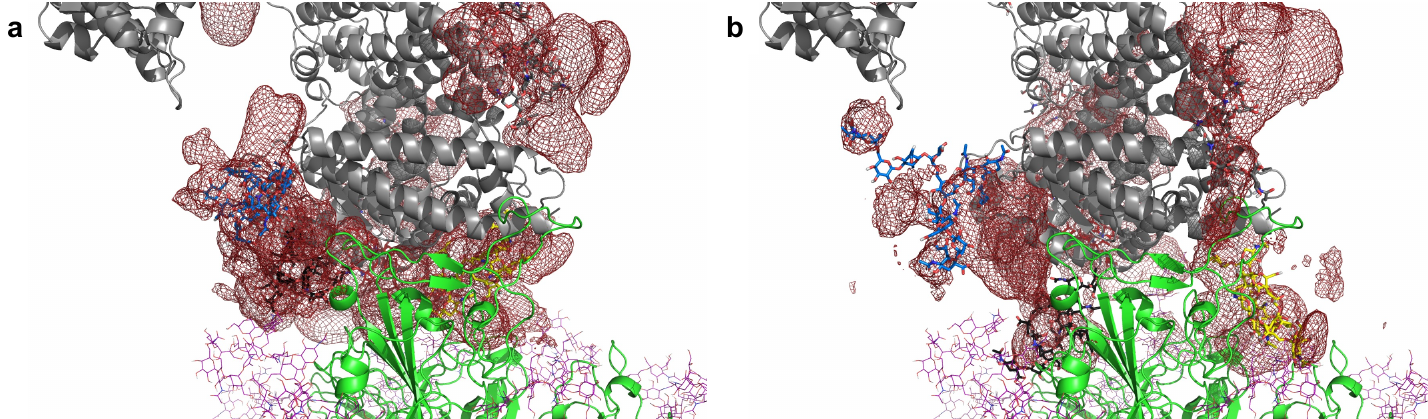

**Figure S18. Average location density maps of sialic acids on ACE2 glycans. (a)** The density map (red mesh) of sialic acids at the termini of the glycans at N53, N90, N322 and N546 as observed in the simulations of unbound ACE2 are superposed onto the ACE2 – Spike complex. **(b)** The density map of the same glycans, as observed in the simulation of the ACE2 – Spike complex. ACE2 in grey, Spike in green. Single, randomly selected conformations of the glycans are shown in blue (N53), yellow (N90), black (N322) and red (N546).

**Supplementary Video**

**Supplementary Video 1. Molecular dynamics simulation of trimeric Spike bound to ACE2.**

The movie highlights a 3 nano-second time segment of the molecular dynamics simulation (from 25 to 28 ns). Trimeric Spike is shown in green, RBD in dark green, and human ACE2 in grey. Complex glycosylation is shown in magenta, Man5 N-glycans in light blue and Man9 N-glycans in orange. Glycans of ACE2 at N53, N90, N322 and N546 are shown in blue, yellow, black and red, respectively.

1. Stawiski, E. W., Diwanji, D., Suryamohan, K., Gupta, R., Fellouse, F. A., Sathirapongsasuti, F., Liu, J., Jiang, Y., Ratan, A., Mis, M., Santhosh, D., Somasekar, S., Mohan, S., Phalke, S., Kuriakose, B., Antony, A., Junutula, J. R., Schuster, S. C., Jura, N., & Seshagiri, S. (2020). Human ACE2 receptor polymorphisms predict SARS-CoV-2 susceptibility. *BioRxiv*. doi:10.1101/2020.04.07.024752
